# PrimalScheme: open-source community resources for low-cost viral genome sequencing

**DOI:** 10.1101/2024.12.20.629611

**Authors:** Chris Kent, Andrew D Smith, John Tyson, Dominika Stepniak, Eddy Kinganda-Lusamaki, Tracy Lee, Mia Weaver, Natalie Sparks, Thomas Brier, Lauren Landsdowne, Sam Wilkinson, Rachel Colquhoun, Áine O’Toole, Placide Mbala-Kingebeni, Ian Goodfellow, Andrew Rambaut, Nicholas Loman, Joshua Quick

**Affiliations:** Institute of Microbiology and Infection, University of Birmingham, Birmingham, UK; British Columbia Centre for Disease Control Public Health Laboratory, Vancouver, Canada; Department of Pathology & Laboratory Medicine, University of British Columbia, Vancouver, Canada; Institut National de Recherche Biomédicale (INRB), Kinshasa, DRC; Service de Microbiologie, Département de Biologie Médicale, Faculté de Médecine, Université de Kinshasa (UNIKIN), Kinshasa, DRC; TRANSVIHMI, Institut de Recherche pour le Développement, Inserm, Université de Montpellier, Montpellier, France; Institute of Ecology and Evolution, University of Edinburgh, Edinburgh UK; Division of Virology, Department of Pathology, University of Cambridge, Cambridge, UK

## Abstract

Viral genome sequencing using the ARTIC protocol has been a vital tool for understanding the spread of epidemics including Ebola, Zika, Covid-19 and Mpox and has seen widespread adoption due to its low cost and high sensitivity. Here, we describe PrimalScheme, an open-source toolkit and website that allows users to easily design primer schemes for amplicon sequencing of viruses, and has generated over 67,000 primer schemes for the global community since 2017. In January 2020, PrimalScheme was used to rapidly generate a primer scheme for SARS-CoV-2, with primer pools distributed to researchers from 44 countries to help scale-up genomic surveillance efforts. Overall, these primers were used to generate an estimated 18M genome sequences and the protocols were viewed online ∼250K times. To complement PrimalScheme, we have built PrimalScheme Labs, a scheme repository which allows users to find and share primer schemes as well as establishing a set of data standards. Through improvements to the primer design process, including the use of discrete primer clouds, we have expanded the use of amplicon sequencing to include diverse virus species. We demonstrate the utility of this approach through a high diversity pan-genotype Measles virus (MeV) scheme. We also demonstrate its use on a high sensitivity, short amplicon Monkeypox virus (MPXV) scheme with over 1000 primers, showing high genome recovery on low-titre clinical samples. These developments have implications for sequencing from samples such as wastewater, for genomic surveillance of endemic pathogens and in preparing for future pandemics.

## Introduction

The rapid and accurate identification of viral pathogens is crucial for effective outbreak management and public health response. Amplicon sequencing generates whole-genome coverage using a tiling approach; it is highly sensitive, low-cost, and compatible with a wide range of clinical sample types. Real-time genomic surveillance using the MinION platform from Oxford Nanopore Technology (ONT) sequencing was first used during the 2015 West African Ebola epidemic, allowing unprecedented responsiveness and resolution into the dynamics of viral outbreaks^1^. We developed PrimalScheme in 2017, where it played a crucial role in generating genome sequences from low-titre Zika virus clinical samples^2^. PrimalScheme is a web-based primer design tool which enables users to design multiplex primer schemes to amplify viral genomes in overlapping amplicons ^3^. At the same time, we published protocols for sequencing amplicon products both for ONT and Illumina. This approach, now commonly referred to as the ‘ARTIC protocol’, is an important method which has been used in response to major outbreaks including Mpox^4,5^, Ebola^1^, and Covid-19^6^.

In January 2020 the first SARS-CoV-2 (then ncov-2019) genome sequence was released^7,8^, and within two weeks we released a primer scheme and sequencing protocol^9,10^. We communicated the initial release and subsequent updates via websites (https://artic.network/ncov-2019) and social media. We also distributed pre-made primer pools at no cost to early adopters who requested them. As demand increased, we collaborated with industry partners (IDT, Sigma-Aldrich) with the capability to mass produce and distribute primer pools to customers. The low cost and availability of ARTIC primers (6p per reaction between April 2020 and March 2021^11^) exhibited a downward pressure on the cost of SARS-CoV-2 sequencing for the duration of the pandemic. Our protocol development focussed on optimising the native barcoding workflow (ONT) while a mixture of academic and commercial groups produced protocols which use PrimalScheme or ARTIC primer designs including; Midnight^12^, COVIDSeq (Illumina), NEBNext ARTIC SARS-CoV-2 (NEB), and QIASeq SARS-CoV-2 (Qiagen). COVIDSeq was the first sequencing-based diagnostic test to be granted emergency use authorisation by the FDA in June 2020^13^.

A recurring problem we faced during the SARS-CoV-2 pandemic was new mutations in primer sites, resulting in reduced amplification efficiency or failure. RNA viruses typically evolve at a rate of ∼2 mutations per month, but in some cases, it is much higher, for example, the Omicon variant contained around 80 new mutations^14^. Primer binding sites cover around 20% of the genome in the ARTIC SARS-CoV-2 schemes, which can be reduced by using longer amplicons e.g. 5% for 1 kb amplicons or 2% for 2 kb amplicons^12,15,16^. Longer schemes are less susceptible to disruption by new mutations, however, as they rely on longer template molecules, they have lower sensitivity. We produced updates to the primer scheme in response to new mutations, which led to different versions being in use at the same time. Over time we developed a semantic versioning system that recognises that some changes have less impact than others, as well using the patch number to indicate balancing. We also produced a set of standards for amplicon sequencing to improve reproducibility. To encourage the uptake of community developed primer schemes, we created a searchable repository called PrimalScheme Labs (labs.primalscheme.com). This allows users to upload the schemes they have developed, making it easier for others to discover them while ensuring interoperability.

The ARTIC protocol has proved to be a robust approach for sequencing during outbreaks but improvements to the design process could allow it to be used more widely. Using a tiling scheme design relies on the representation of all amplicons to generate an intact consensus sequence. It has been observed that primer interactions reduce the amplification efficiency of certain amplicons resulting in dropouts. The cause was shown to be 3’ complementarity between primers leading to dimer formation^17^. Interactions are an important factor limiting the complexity of multiplex PCR reactions as the number of possible interactions increases exponentially as primers are added^18–22^. Primer schemes for the large 200 kb DNA genome of MPXV have used long amplicons e.g. 2 kb to reduce the number of amplicons^4^. This limits the use of long amplicons for low titre or degraded samples such as wastewater. A significant challenge has been developing primer schemes tolerant of sequence diversity which are desirable for surveillance and pandemic preparedness^23,24^. Previous attempts to develop a pan-genotypic scheme for MeV with 12% diversity^25^ by selecting primer pairs in conserved regions of the genome were unsuccessful. When sequence diversity exceeded ∼5% there were too few conserved positions to generate a continuous tiling path across the genome while the use of degenerate primers results in primers with a melting temperature (Tm) outside the target range (**Supplementary Figure 4**). The latest version of PrimalScheme has significant improvements which will enable the surveillance of endemic viruses and larger, higher complexity schemes greatly increasing the power of the method.

## Methods

### Determining usage

Log files from the PrimalScheme website up to 28 November 2024 were parsed to determine the number of jobs processed. Primer shipping information was taken from a spreadsheet used to record primer shipments. Primers were shipped as 100 µM pools at ambient temperature. Total views for the SARS-CoV-2 sequencing protocol were taken from the metrics tab on the protocol page on 9 December 2024. The usage of primer schemes to generate genome sequences for SARs-CoV-2 was calculated from **Supplementary Table S1** of Hunt *et al*., 2024.

### primalscheme3

primalscheme3 (https://github.com/artic-network/primalscheme3) is the latest version of PrimalScheme, written in Python 3 and Rust, and uses primer3^26^ for thermodynamic calculations. The steps are as follows:

- **Primer digestion:** The input multiple sequence alignment (MSA) is stored as a 2D array, with genomes as rows and bases as columns. Primer clouds are generated by selecting a fixed 3’ index, and recursively extending the primer in the 5’ direction until the desired Tm is achieved (59 - 62°C) (**Figure 1**), producing a separate primer for each genome. The occurrence of each primer with the cloud is calculated and optionally filtered based on a user-specified threshold. Each primer undergoes checks for GC%, hairpin formation, sequence complexity and self dimerization, and clouds that contain failing primers are discarded.
- **Primer pair generation:** For each forward primer, all reverse primers which would form an amplicon within the required size range, are checked for interactions and mispriming.
- **Pool generation**: Depending on the run mode one of two solving algorithms is run. Scheme is for tiling amplicon schemes while panel is used for discontinuous regions.
  - **Scheme:** The first valid primer pair from the left is selected and added to the pool to become the leading primer pair. All primer pairs that overlap with the leading primer pair are scored and sorted on: the size of overlap, number of primers and amplicon size (**Supplementary Figure 2**). Each overlapping primer pair is checked to see if it can be added to any pool other than the leading primer pair’s pool. If a valid primer pair is found, it is added, it becomes the leading primer pair, and the process repeats. If not, the solver attempts to use the backtrack logic (**Supplementary Methods**). If no overlapping primer pairs can be added, a gap is unavoidable, and a walking primer pair is added by iterating over all primer pairs further along the genome. The first primer pair that can be added to any pool is selected and set as the leading primer pair for the next stage.
  - **Panel:** A region file specifies the region and its score, which is passed into a score array, with a specified score for each genomic position. With primer pairs scored by the sum of the covered genome positions, and sorted in descending order, until a valid primer pair is found and can be added. The score of the region covered by the primer pair is set to 0, and all primer pairs are then rescored and resorted, and the process repeats.
- **Figure creation:** Interactive plots are produced, displaying information about the scheme and input alignment (**Supplementary Figure 3**).

**Figure 1.**
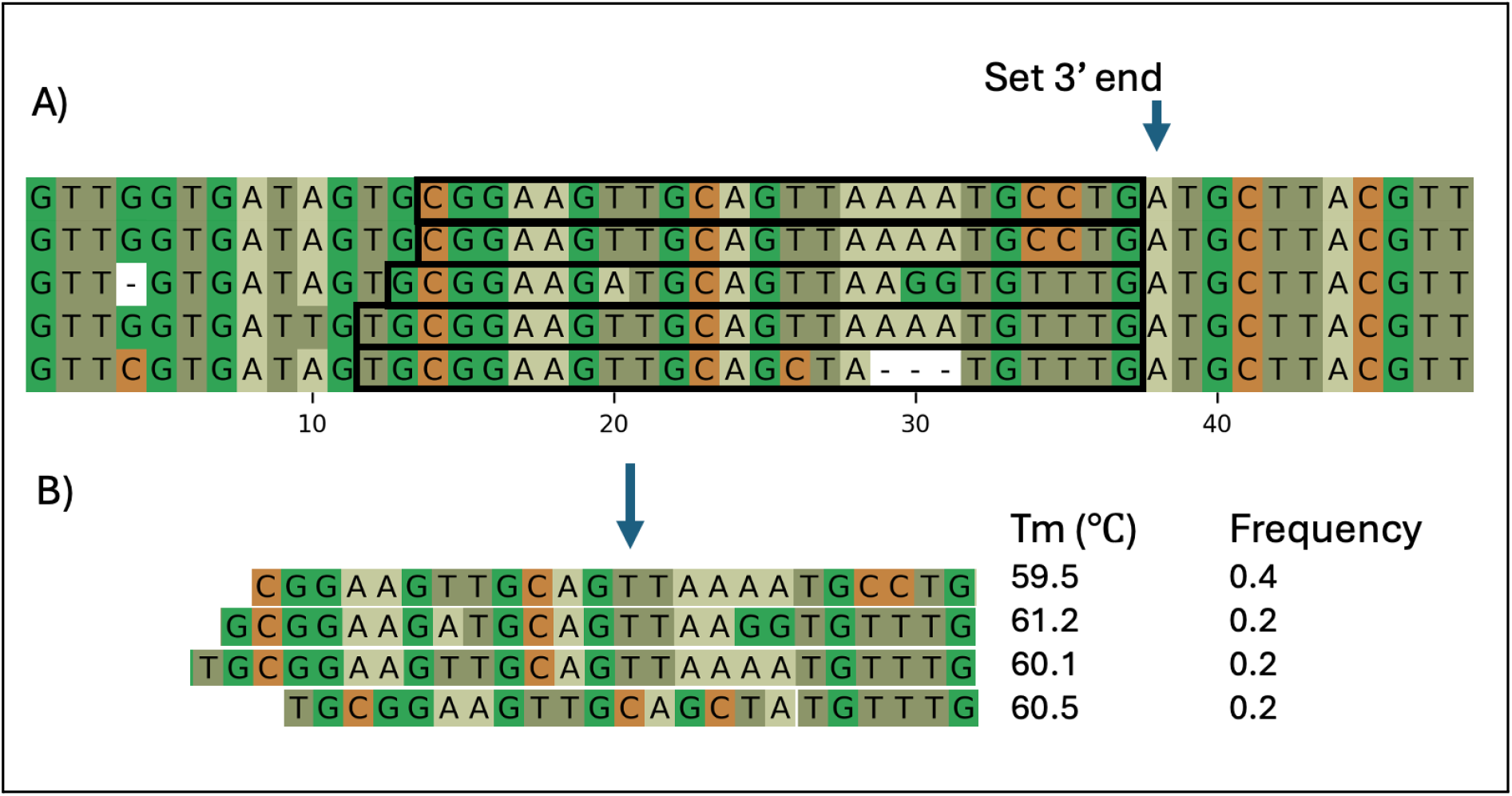
An overview of the digestion algorithm which generates the discrete primer clouds. **A)** shows an example input alignment. After establishing the 3’ end, primers are extended until reaching the target Tm range (black boxes). **B)** shows the resulting primer cloud, Tm of each primer, and the frequency each primer occurs in the original alignment.

### MeV scheme design

A total of 895 full-length MeV genomes were downloaded from NCBI Virus and aligned with MAFFT v7.490^27^. A tree was generated using FastTree using run mode ‘`--fastest`’^28^, and then phylogenetically downsampled with Treemmer to 0.95 Relative Tree Length (RTL) to leave 376 genomes^29^. The downsampled genomes were aligned with MAFFT with the primary reference first in the file. The primary reference is the reference genome that the coordinate system the primer BED file is based on. The alignment was used as the input to PrimalScheme v3.0.0 with the command `’primalscheme3 scheme-create --min-base-freq 0.01 --high-gc –backtrack --amplicon-size 400’.`

### Measles virus (MeV) sequencing

Oligos were ordered from Integrated DNA Technologies (IDT) in 96-well plate format normalised to 10 nmole yield. A script `bed2idt` (https://github.com/artic-network/bed2idt) was used to parse the sequences into a bulk input spreadsheet. Plates were briefly spun down and oligos were resuspended to 100 µM with Tris-EDTA (TE) buffer. Oligos were left to rehydrate for 30 minutes and pipette mixed 10 times. 5 µl of each primer was combined for each pool, to create the 100 µM equimolar stocks for each pool using an OpenTron OT2 and `primalpooling` protocols (https://github.com/artic-network/primalpooling).

RNA from 8 MeV strains (Genotypes A, B3, D4 and D5) (**Supplementary Table 1**) from the National Collection of Pathogenic Viruses (NCPV) were diluted 1:100 in nuclease-free water. cDNA was generated by combining 8 µL of diluted RNA and 2 µL of LunaScript RTSuperMix (5X) (NEB) and incubating at 25°C for 2 minutes, 55°C for 10 minutes and 95°C for 1 minute. Multiplex PCR was performed by combining 12.5 µL Q5 Hot Start High Fidelity 2X Master Mix (NEB), 0.73 µL primer pool 1 or 2 (15 nM each primer), 2.5 µL of cDNA mixture from the previous step and 9.27 µL nuclease-free water. The ARTIC LoCost protocol v4^30^ was followed from step 7 onwards with no other changes. The final sequencing library (25 fmol) was loaded onto an R10.4.1 flow cell (ONT) and sequenced on a GridION for 48 hours. Basecalling was performed on board using the HAC model and ‘Barcode both ends’ option enabled. Consensus genomes were generated using fieldbioinformatics 1.2.4 (https://github.com/artic-network/fieldbioinformatics) with the command `’artic minion –medaka` `--normalise 400 --threads 4 --medaka-model r1041_e82_400bps_hac_v4.3.0’.`

### MPXV scheme design

MPXV genomes were downloaded from online databases and then deduplicated to 306 genomes. Of the 306 genomes, 76 were Clade I and 230 were Clade II. Clade I genomes were aligned to reference NC_003310 and clade II genomes were aligned to reference NC_063383 using squirrel (https://github.com/artic-network/squirrel-nf). These were then consensus-aligned to each other using MAFFT. For all genomes, the 3’ inverted terminal repeat (ITR) is masked with Ns as it is identical to the 5’ ITR. The primary reference was KJ642613 commonly used as a Clade I reference genome. PrimalScheme v3.0.0 was run using the command `’primalscheme3 scheme-create` `--ignore-n --amplicon-size 400 --min-base-freq 0.025’`. The scheme was remapped to the Clade IIb reference genome (**Supplementary Table 2**) using ‘`primalbedtools remap’` to perform coverage analysis of Clade IIb samples. The scheme was published to PrimalScheme Labs (https://labs.primalscheme.com) using `’primal-page create’` (https://github.com/artic-network/primal-page).

### MPXV sequencing

Two extractions of clinical samples were selected for sequencing which had previously tested positive for Mpox with Ct values of 26.1 and 31.0. Multiplex PCR was performed by combining 5 µL 5X Q5 Reaction Buffer, and 0.25 µL Q5 Hot Start DNA polymerase (NEB), 2.2 µL primer pool 1 or 2 (100 µM), 0.5 µl 10 mM dNTPs, 2.5 µL of DNA and 14.55 µL nuclease-free water and cycling conditions of 98°C for 30 seconds for 1 cycle followed by 98°C for 15 seconds and 65°C for 5 minutes for a further 35 cycles. Amplicons were prepared for sequencing using a modified Illumina DNAPrep workflow^31^. Libraries were sequenced on a MiSeq i100 (Illumina) using a 25M 300-cycle kit. Primary sequence processing and analysis were performed using the ncov2019-artic-nf pipeline (https://github.com/jts/ncov2019-artic-nf) using the NC_063383 reference genome and the Clade IIb remapped BED file, and coverage data was extracted using ncov-tools (https://github.com/jts/ncov-tools).

## Results

### Open-source resources for viral genome sequencing

There have been three major iterations of the primalscheme.com website (https://github.com/artic-network/primalweb). The initial site was available from April 2017 and was updated in June 2020 and November 2024. In this time the website has processed ∼67.7K jobs, with usage increasing over time (**Figure 2**). The spike in usage in August 2022 was a result of jobs being submitted directly via the website API. To improve visibility and reuse of primer schemes, we created PrimalScheme Labs, which allows users to browse and search for primer schemes. New schemes can be added using `primal-page` to generate the required format to create a pull request. There are currently 49 schemes in the repository including 16 submitted by community members.

**Figure 2.**
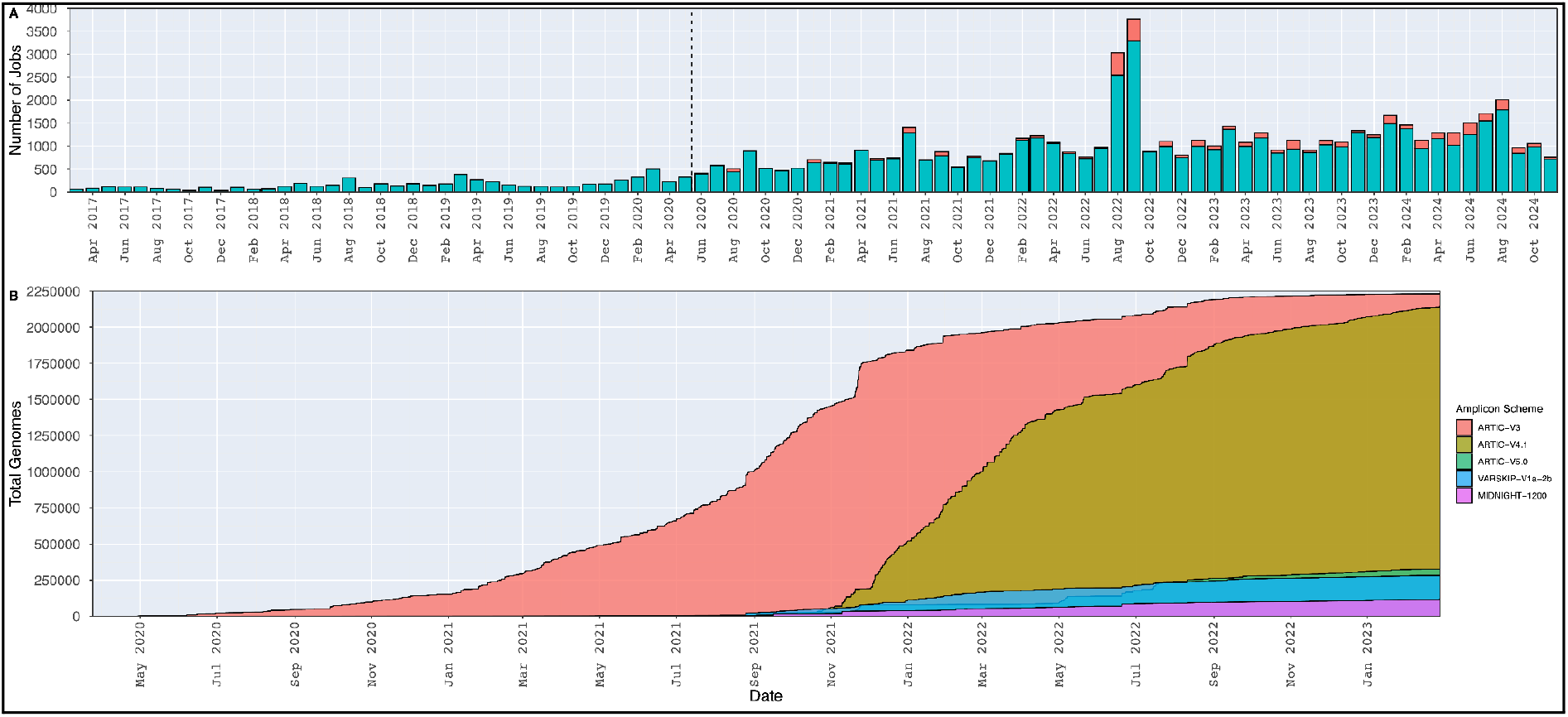
**A)** Number of jobs processed by primalscheme.com by month. Stratified by completed (green) and failed (red) where available. The vertical dashed line shows when an updated version of the website was released. **B)** The cumulative sum of genome sequences deposited in ENA by identified primer scheme.

### ARTIC resources for SARS-CoV-2

The first version of the ARTIC SARS-CoV-2 sequencing protocol was published on 22nd January 2020 and over the next three months, we shipped primer pools to 128 users in 44 different countries supporting the establishment of sequencing capacity in these laboratories. The protocol received updates in April 2020, August 2020 and June 2024 to improve double barcoding detection and reduce costs^32^. The combined protocol versions received 247,463 page views. The first stable version of the primer scheme was V3 which was released in March 2020. This was succeeded by V4 in June 2021 due to mutations in the Spike protein associated with Beta/Gamma/Delta variants, V4.1 in January 2022 following the emergence of the Omicron variant and V5.3.2 in January 2023 in response to the large number of Omicron sublineages circulating at that time (https://github.com/artic-network/primer-schemes). Of the 5.29M SARS-CoV-2 genomes uploaded to ENA, 4.71M (89%) were identified to have been generated using an ARTIC scheme (V3, V4.X or V5.X.X) and another 0.25M (4.7%) generated using other schemes designed using PrimalScheme (Midnight, VarSkip)^33^. Assuming a similar usage pattern for the sequences submitted to GISAID, we estimate over 18M genomes have been generated using these schemes.

### Data standards for ARTIC sequencing protocols

Performing downstream analysis appropriately is vitally important to consensus genome accuracy. To improve reproducibility we developed a schema including semantic versioning and specifications of essential file formats. Each scheme has a unique ID created by combining the key information about the scheme `{scheme name}/{amplicon size}/{scheme version}.` The `{scheme name}` is a text field describing the scheme, which often includes the project or target organism e.g. `artic-sars-cov-2`. The `{amplicon size}` is the amplicon length for the scheme. The `{scheme version}` is the iteration of the scheme in the form `v{x}.{y}.{z}-{clade},` with `{x}` denoting a major redesign or new scheme, `{y}` denoting a change to the primers and `{z}` denoting a change to the primer balancing. While a single primer scheme can amplify genomes from different clades or genotypes, we have found it is still preferable to use a closely related reference genome for consensus generation. To allow this, the optional `{clade}` denotes if the scheme has been remapped to a different primary reference. Each scheme requires two essential files; the `reference.fasta` containing the sequence of the primary reference, and the `primer.bed` containing the sequence and location of the primers. We have created a specification for both input files which are compatible with the fieldbioinformatics pipeline (https://github.com/artic-network/fieldbioinformatics).

### Pan-genotypic amplicon sequencing of MeV

The pan-genotypic MeV scheme (https://labs.primalscheme.com/detail/artic-measles/400/v1.0.0/) consisted of 46 amplicons and an average of 4 primers per cloud. The majority of primers had an exact match to an input genome with an average of 0.05 mismatches per primer. The scheme covers the entire coding region of the genome missing only the first 26 and last 37 bp relative to the primary reference NC_001498.1. Sequencing of 8 MeV strains from a viral culture collection showed all amplicons had at least 20x depth of coverage (**Figure 3)**. Of the strains sequenced, 5 were genotype A while the others were B3, D4 and D5 according to the strain information (**Supplementary Table 1**). Three of the strains; Schwartz (0809212v), Schwartz (0007233v) and Moraten, were found to have identical or near-identical reference genomes which had been included in the input dataset for the scheme design (**Supplementary Figure 6**). These are vaccine strains derived from the Edmonston strain isolated in 1954. There were no reference sequences for the B3, D4 or D5 strains identified but in each case, there were examples of the genotype in the input dataset. For the M/F intergenic region, there was amplicon coverage but a subset of the amplicons contained variable length deletions resulting in low coverage for some positions. We confirmed these deletions using metagenomic sequencing (data not shown) and believe it to be a genuine variation linked to the passage of the virus^34^ and not related to the sequencing approach.

**Figure 3.**
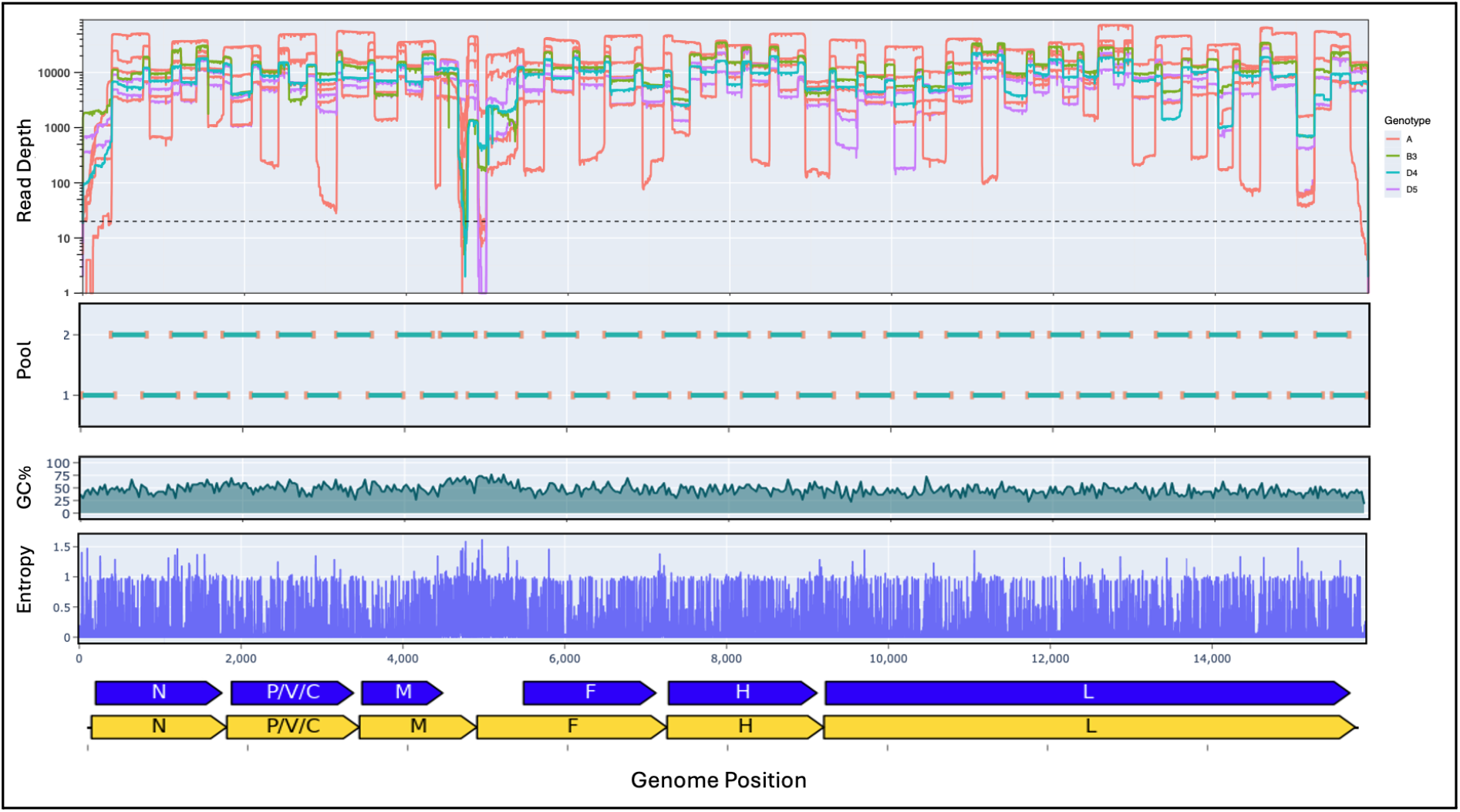
Scheme design and sequencing coverage of MeV primer scheme. The top panel shows the read depth across the MeV genome for each genotype, A in red, B3 in green, D4 in blue, and D5 in purple. The second, third and fourth panel shows the positions of the amplicons, GC content and Shannon entropy across the genome. The fifth panel shows genes (yellow) and the CDS (blue). All features are relative to the coordinates of reference genome NC_001498.

### High sensitivity amplicon sequencing of MPXV

The high sensitivity MPXV scheme consisted of 553 ∼400 bp amplicons covering 97.1% of the 3’ UTR masked reference genome. The scheme had an average of 1.07 primers per cloud, which was consistent with the maximum pairwise distance of 0.4% between sequences, calculated using MEGA^35^. When comparing primers to the reference genomes for each clade the average mismatches per primer was 0.003. The scheme was tested on two clinical samples of Ct 26.1 and Ct 31.0, representing medium and low viral loads (**Figure 4**). There was at least 10x depth of coverage for 98.0% of the genome for the medium viral load sample and 92.5% of the genome for the low viral load sample. Of the 553 amplicons, only four were below the coverage threshold of 10x in the medium viral load sample; 55, 362, 453 and 454. Three of these can be explained due to differences between the Clade I and II reference genomes. One of the primers for amplicon 55 is not present in the Clade IIb reference genome due to a relative deletion. An insertion in Clade IIb relative to Clade Ia within the amplicon 453 and 454 overlap region results in two amplicons ∼2,800 bp in length which prevents them from amplifying efficiently.

**Figure 4.**
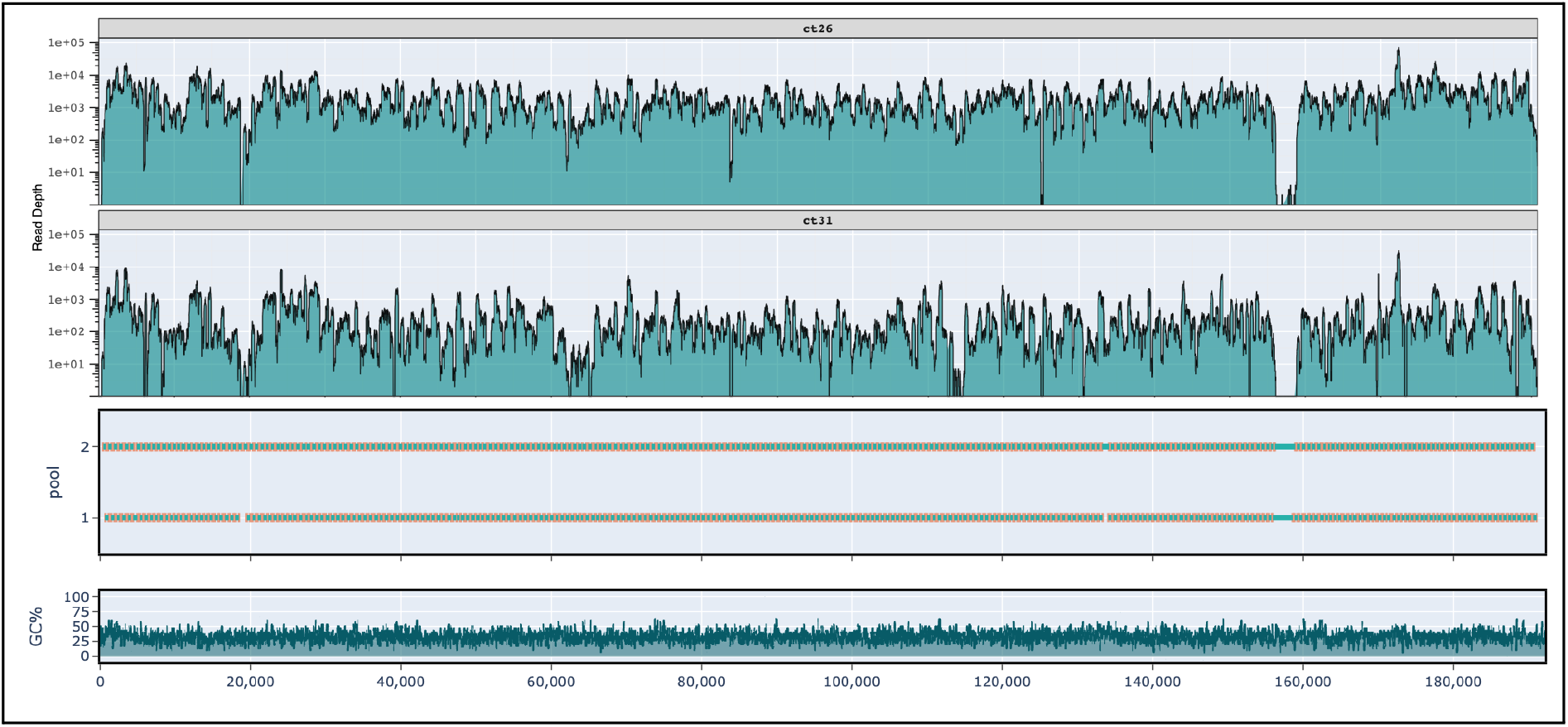
Depth of coverage for the MPXV 400 bp scheme for medium (first panel) and low (second panel) viral load samples. Positions of amplicons (third panel) and GC content (fourth panel) are shown below. All coordinates are relative to the Clade IIb reference genome NC_063383 with the 3’ UTR masked.

## Discussion

### Open-source resources for viral genome sequencing

Open-source resources democratise access to genomic surveillance, particularly benefiting low and middle-income countries (LMICs) that disproportionately bear the human cost of infectious diseases. They provide opportunities for local scientists to develop expertise, build local capacity, and reduce dependency on proprietary reagents and technologies. ARTIC open-source primer schemes, protocols and tools are important community resources for pathogen surveillance as demonstrated by the usage metrics. Analysis of data deposited in public databases, showed that an estimated 18M SARS-CoV-2 genomes used these primer schemes. Surveillance networks are more effective when data is generated over the widest possible area, with low-cost, accessible sequencing approaches enabling the largest network of laboratories to contribute. The collaborative nature of open-source resources allows for quick dissemination of new methods during outbreaks. Leveraging online platforms like GitHub.com and protocols.io, we were able to publish the SARS-CoV-2 sequencing protocols less than a week after the first genome sequence was available. The rapid sharing of the first genome sequence was a critically important step in the development of diagnostic and sequencing assays. Open-source protocols allow improvements to be suggested by the community. For example, analysis of dimers generated by ARTIC V3 primers by researchers in Japan led to improvements in PrimalScheme software and the quality of future scheme designs. Other groups took the materials provided under the permissive CC-BY-4.0 license and customised them for their own needs. This led to the development of important products such as NEBNext ARTIC SARS-CoV-2 (NEB) and CovidSeq (Illumina). Transparent processes are essential to establish trust in the quality of surveillance data. Every stage of the sequencing pipeline from primer design software to sequencing analysis pipelines is available online. This provides users with the ability to replicate work and find problems, examples of this include the discovery of artefacts in consensus genome sequences caused by data quality issues or improper analysis approaches^36^. We have established a set of data standards for amplicon sequencing to try to promote reproducibility and interoperability with analysis pipelines going forward.

### Primer schemes for diverse virus species

Our approach to primer design addresses an important question in amplicon sequencing; how do you capture more diversity without sacrificing specificity? A primer scheme using discrete primer clouds was able to completely sequence different genotypes of MeV without modifying the PCR conditions to reduce the stringency. Single primer pairs targeting conserved regions of the genome^37^ works well for small numbers of closely related input genomes, but is not scalable. We were able to use ∼900 publicly available MeV genomes in this scheme design, incorporating phylogenetic downsampling to capture as many different clades as possible. The minimum base frequency argument in primalscheme3 provides additional control over the size of the primer clouds by removing the least common primers. The MeV scheme used 4 primers per cloud but we have successfully tested schemes with up to 40 primers per cloud (data not shown). Further work is needed to predict the effect of mismatches within primers, which will allow us to optimise the size of the primer clouds. A scheme we designed for another study was able to generate full genomes from 12 different genotypes of Hepatitis B virus (HBV), with an average cloud size of 8.7 primers. HBV is known to be challenging to sequence given the high diversity and low viral loads encountered^38^, demonstrating the potential of this approach.

### Increasing the complexity of multiplex PCR reactions

We also demonstrated a high sensitivity MPXV scheme using 400 bp amplicons as an alternative to previous schemes which have ∼2kb. This increase in sensitivity offers opportunities to perform wastewater and environmental surveillance where titres are expected to be low. Primer interactions have limited the scale of multiplex PCR reactions and a variety of different approaches to avoid them have been used including; digestible primers (AmpliSeq), microfluidic devices (Fluidigm) or droplets (Raindance). Our thermodynamic approach can predict interactions with high sensitivity and specificity which allows more complex reactions to be designed. Simulations have shown that schemes containing thousands of primers are feasible (**Supplementary Figure 5**), paving the way for bacterial whole-genome schemes and multi-pathogen panels in future.

### Effect on cost and time

One key advantage of amplicon sequencing as a surveillance approach is the low cost of around £10 per sample^32^. The use of primer clouds or larger schemes increases the cost however the effect is modest as oligos account for a small amount of the overall sequencing cost. Resuspending and pooling more oligos manually, takes longer but this is a one-time upfront task. We have found it easier to order these schemes in 96-well plates and resuspend them using an OpenTrons liquid handling robot. This way it is possible to resuspend and pool oligos in a few hours of mostly hands-off time. Having multiple forward and reverse primers complicates the rebalancing of amplicons based on coverage. We have observed that using equimolar pooling gives behaviour similar to that of a conventional scheme but further work is needed to determine the best rebalancing strategy.

### Future perspectives

The high sensitivity MPXV scheme highlighted some difficulties in working with pan-clade schemes. When using a single primer scheme to amplify genomes of different clades, we found a closely related reference is still preferable for reference based consensus generation. MPXV has large clade specific insertions and deletions which makes it impossible to remap all primers from one reference to another exactly. Remapping the Clade Ia reference to Clade IIb resulted in the loss of one of the primer sites whilst another two moved, resulting in the amplification failing. Further work is needed to investigate designing primers using a genome graph which would enable multiple paths to be created. It may also be possible to create an end-to-end process where consensus genomes are also generated using this approach.

This analysis shows how valuable ARTIC open-source resources are for viral genome sequencing especially in LMICs where they have proven cost-effective. These advances in primer design for viral genome sequencing will allow the surveillance of endemic diseases, as well as improving our ability to react to outbreaks and threats.

## Author contributions (CRediT taxonomy)

Conceptualization - CK, AS and JQ; Methodology - CK, AS, JT and JQ; Software - CK and AS;

Formal Analysis - CK, JT and JQ;

Investigation - CK, AS, JT, DS, EL, TL, MW, LL, SW, RC, AO and JQ;

Resources - JT, EL, PM, IG, AR, NL and JQ; Writing – Original Draft, CK, JQ;

Writing – Review & Editing, CK, JT, EL, TB, NS, LL, PM, IG, AR, NL and JQ; Visualization - CK and JT;

Supervision - JT, AR, NL and JQ;

Funding Acquisition - PM, IG, AR, NJL and JQ

## Acknowledgements

We would like to thank Dr. Barry Atkinson and the National Collection of Pathogenic Viruses (NCPV) for providing the MeV RNA. The PrimalScheme website is hosted on CLIMB-BIG-DATA. The PrimalScheme logo was designed by Oliver Pybus. This work was supported by the Wellcome Trust (Collaborative Award 206298/Z/17/Z – ARTIC). The Mpox scheme was developed by the Modjadji project funded by the Bill & Melinda Gates Foundation and sequencing testing performed at the BCCDC PHL. JQ holds a UKRI Future Leaders Fellowship.

## Data availability

The data are available under ENA Project PRJEB81871.

## References

1. Quick, J. et al. Real-time, portable genome sequencing for Ebola surveillance. Nature 530, 228–232 (2016).

2. Faria, N. R. et al. Establishment and cryptic transmission of Zika virus in Brazil and the Americas. Nature 546, 406–410 (2017).

3. Quick, J. et al. Multiplex PCR method for MinION and Illumina sequencing of Zika and other virus genomes directly from clinical samples. Nat. Protoc. 12, 1261–1276 (2017).

4. Chen, N. F. G. et al. Development of an amplicon-based sequencing approach in response to the global emergence of mpox. PLOS Biol. 21, e3002151 (2023).

5. Isabel, S. et al. Targeted amplification-based whole genome sequencing of Monkeypox virus in clinical specimens. Microbiol. Spectr. 12, (2024).

6. Charre, C. et al. Evaluation of NGS-based approaches for SARS-CoV-2 whole genome characterisation. Virus Evol. 6, veaa075 (2020).

7. Wu, F. et al. Severe acute respiratory syndrome coronavirus 2 isolate Wuhan-Hu-1, complete genome. (2020).

8. Wu, F. et al. A new coronavirus associated with human respiratory disease in China. Nature 579, 265–269 (2020).

9. ARTIC Network. ARTIC have developed a set of lab and bioinformatics protocols for the nCoV-2019 virus using a targeted multiplex primer scheme. This package enables direct sequencing and real-time analysis with RAMPART of the coronavirus on nanopore sequencers: https://artic.network/ncov-2019 https://t.co/hPbvmpX5DE. Twitter https://x.com/NetworkArtic/status/1220673867473092609 (2020).

10. Quick, J. nCoV-2019 sequencing protocol v1. Preprint at 10.17504/protocols.io.bbmuik6w (2020).

11. ARTIC Network. nCoV-2019 Version 3 Amplicon Release. https://artic.network/resources/ncov/ncov-amplicon-v3.pdf (2020).

12. Freed, N. E., Vlková, M., Faisal, M. B. & Silander, O. K. Rapid and inexpensive whole-genome sequencing of SARS-CoV-2 using 1200 bp tiled amplicons and Oxford Nanopore Rapid Barcoding. Biol. Methods Protoc. 5, bpaa014 (2020).

13. FDA. Coronavirus (COVID-19) Update: FDA Authorizes First Next Generation Sequence Test for Diagnosing COVID-19. FDA https://www.fda.gov/news-events/press-announcements/coronavirus-covid-19-update-fda-authorizes-first-next-generation-sequence-test-diagnosing-covid-19 (2020).

14. Markov, P. V. et al. The evolution of SARS-CoV-2. Nat. Rev. Microbiol. 21, 361–379 (2023).

15. Kuchinski, K. S. et al. Mutations in emerging variant of concern lineages disrupt genomic sequencing of SARS-CoV-2 clinical specimens. Int. J. Infect. Dis. 114, 51–54 (2022).

16. Resende, P. C. et al. SARS-CoV-2 genomes recovered by long amplicon tiling multiplex approach using nanopore sequencing and applicable to other sequencing platforms. Preprint at 10.1101/2020.04.30.069039 (2020).

17. Itokawa, K., Sekizuka, T., Hashino, M., Tanaka, R. & Kuroda, M. Disentangling primer interactions improves SARS-CoV-2 genome sequencing by multiplex tiling PCR. PLOS ONE 15, e0239403 (2020).

18. Fuchs, J. et al. varVAMP: automated pan-specific primer design for tiled full genome sequencing and qPCR of highly diverse viral pathogens. 2024.05.08.593102 Preprint at 10.1101/2024.05.08.593102 (2024).

19. Johnston, A. D., Lu, J., Ru, K., Korbie, D. & Trau, M. PrimerROC: accurate condition-independent dimer prediction using ROC analysis. Sci. Rep. 9, 209 (2019).

20. Linhart, C. & Shamir, R. The degenerate primer design problem. Bioinforma. Oxf. Engl. 18 Suppl 1, S172–81 (2002).

21. Wang, M. X. et al. Olivar: automated variant aware primer design for multiplex tiled amplicon sequencing of pathogens. bioRxiv 2023.02.11.528155 (2023) doi:10.1101/2023.02.11.528155.

22. Xie, N. G. et al. Designing highly multiplex PCR primer sets with Simulated Annealing Design using Dimer Likelihood Estimation (SADDLE). Nat. Commun. 13, 1881 (2022).

23. Lambisia, A. W. et al. Genomic epidemiology of human adenovirus F40 and F41 in coastal Kenya: A retrospective hospital-based surveillance study (2013–2022). Virus Evol. 9, vead023 (2023).

24. Maes, M. et al. Multiplex MinION sequencing suggests enteric adenovirus F41 genetic diversity comparable to pre-COVID-19 era. Microb. Genomics 9, (2023).

25. Bellini, W. Genetic Diversity of Wild-Type Measles Viruses: Implications for Global Measles Elimination Programs. Emerg. Infect. Dis. 4, 29–35 (1998).

26. Untergasser, A. et al. Primer3—new capabilities and interfaces. Nucleic Acids Res. 40, e115–e115 (2012).

27. Katoh, K. MAFFT: a novel method for rapid multiple sequence alignment based on fast Fourier transform. Nucleic Acids Res. 30, 3059–3066 (2002).

28. Price, M. N., Dehal, P. S. & Arkin, A. P. FastTree: Computing Large Minimum Evolution Trees with Profiles instead of a Distance Matrix. Mol. Biol. Evol. 26, 1641–1650 (2009).

29. Menardo, F. et al. Treemmer: a tool to reduce large phylogenetic datasets with minimal loss of diversity. BMC Bioinformatics 19, 164 (2018).

30. Quick, J. & Lansdowne, L. ARTIC SARS-CoV-2 sequencing protocol v4 (LSK114) v2. Preprint at 10.17504/protocols.io.bp2l6n26rgqe/v4 (2024).

31. Hickman, R. et al. Rapid, high-throughput, cost-effective whole-genome sequencing of SARS-CoV-2 using a condensed library preparation of the Illumina DNA Prep kit. J. Clin. Microbiol. 62, e00103–22 (2024).

32. Tyson, J. R. et al. Improvements to the ARTIC Multiplex PCR Method for SARS-CoV-2 Genome Sequencing Using Nanopore. http://biorxiv.org/lookup/doi/10.1101/2020.09.04.283077 (2020) doi:10.1101/2020.09.04.283077.

33. Hunt, M. et al. Addressing pandemic-wide systematic errors in the SARS-CoV-2 phylogeny. Preprint at 10.1101/2024.04.29.591666 (2024).

34. Beaty, S. M. & Lee, B. Constraints on the Genetic and Antigenic Variability of Measles Virus. Viruses 8, 109 (2016).

35. Tamura, K., Stecher, G. & Kumar, S. MEGA11: Molecular Evolutionary Genetics Analysis Version 11. Mol. Biol. Evol. 38, 3022–3027 (2021).

36. Wilkinson, S., Groves, N., Quick, J. & Loman, N. J. Erroneous Mutations Associated with 64_L-60_R Primer-Dimer in ARTIC 4/4.1 - Laboratory. ARTIC Real-time Genomic Surveillance https://community.artic.network/t/erroneous-mutations-associated-with-64-l-60-r-primer-dimer-in-artic-4-4-1/419/1 (2022).

37. Quick, J. et al. Multiplex PCR method for MinION and Illumina sequencing of Zika and other virus genomes directly from clinical samples. Nat. Protoc. 12, 1261–1276 (2017).

38. Lumley, S. F. et al. Whole genome sequencing of hepatitis B virus (HBV) using tiled amplicon (HEP-TILE) and probe-based enrichment on Illumina and Nanopore platforms. Preprint at 10.1101/2024.09.11.24313306 (2024).

