## Supplementary Material for "PrimalScheme: open-source community resources for low-cost viral genome sequencing"

### Supplementary Methods

##### primalscheme3

An overview of the scheme design process and primalscheme3 software is shown in **Supplementary Figure 1**. Detailed descriptions for each stage are as provided below.

##### Creating the mispriming db

To detect possible sites where primers could misprime in the target genomes, each input genome is digested into the primer minimum length (default 18 bases), and the sequence is used as the key, with the index and direction stored in a file-based database. Therefore, when a primer is being assessed for mispriming sites, the last N bases from its 3’ end can be queried within the database, to find all locations and directions this primer can bind, in a constant time. To account for mismatches, all possible sequences with 1-mutation from the original query are also searched. Two mispriming sites on opposite strands less than 2x amplicon length would be deemed to produce an off-target product, and would not be allowed.

##### Digestion and primer pair generation

The MSA is read into a numpy 2d array for rapid indexing, with each genome becoming a row in the array, and each base being a column position. Gap bases ‘-’ located on the 3’ and 5’ end of the genomes are replaced with an empty base. A column index is used as the 3’ end of the primer, therefore, forward-kmers and reverse-kmers are digested separately. For each genome in the MSA an initial slice is taken `(msa[genome_index, 3p_index - primer_min_length : 3p_index])` for forward-kmers, and `(msa[genome_index, 3p_index : 3p_index + primer_min_length])` for reverse-kmers, to this slice bases are recursively appended to the 5’ end of the primer until the Tm, calculated by primer3 thermodynamic library [1], is within the required range. Reverse primers are reverse complemented, as the sequence is stored as 5’ to 3 direction’.

The recursive nature of the algorithm allows ambiguous bases in the input genomes to be handled. When an ambiguous base is added to the slice, it is expanded to all base representations, with each base added to a separate slice, to ensure all primers fall in the required Tm range. This provides thermodynamic normalisation of degenerate bases in the input data but can produce a large number of primers for highly degenerate input. The algorithm also allows the handling of gap bases caused by indels, by skipping gap bases (‘-’) and moving to the next base position, and returning errors on non-DNA bases / invalid indexes.

This method produces a primer cloud, containing a sequence with no mismatches to every input genome contained in this region of the MSA, except for edge cases, where non-IUPAC bases are found or the primer has to extend outside the size of the MSA. If a minimum frequency is specified by the user, primer frequencies are calculated using `(1 / (number of primers for genome)) / total genomes`. Use of the `1 / (number of primers for genome`. Therefore, if a genome has degenerate bases and produces a large number of primers, each primer will get a low frequency. Any primer below the frequency is removed. This frequency filtering does remove primer sequences from the cloud, allowing potential mismatches. However, using frequency filters ensures that point mutations or sequencing errors, which are only found in a few genomes, do not make their way into the primers. All remaining primers within the cloud undergo thermodynamic checking, using parameters adapted from Quick et al. [2]. GC content is required to be between 30 and 55%, or 40 and 65% for high-gc mode. Hairpin formation temperature must be less than 47.0 degrees, no homo- and hetero-dimer formation, and tracks of homopolymers less than 5. If any primer within the cloud fails, the entire cloud is discarded.

This primer cloud is represented by a Kmer object, which contains a set start or end position for forward or reverse primers, respectively, and the set of primer sequences. Primer sequences are always displayed in the 5’ to 3’ orientation. The 3’ index of the kmer is remapped from the MSA index to the index of the primary reference (first genome in msa).

**Panel mode**

This version introduced the ability to design amplicons which target specific regions, of a variable number of target genomes. A region.bed file specifies the region and its score, which is passed into a score array that specifies a score for each genomic position. An additional run option can use the Shannon entropy at each position as the score, to target variable regions. The genomes are digested into primer pairs, which are scored by the sum of the covered genome positions. The primer pairs are checked in descending score until a valid primer pair is found and can be added. The score of the region covered by the primer pair is set to 0, and all primer pairs are then rescored and resorted, and the process repeats. To ensure overlapping amplicons are favourable, adjacent edges of the score array are heavily favoured (default 10x score). This progress repeats until either all regions are covered or until the specified number of amplicons are added (**Supplementary Figure 2)**.

**Scheme Mode**

Scheme mode generates a tiling scheme, across variable numbers of MSAs. All the genomes are digested into primer pairs, and for each MSA, the first valid primer pair from the left is selected and added to the pool, to become the leading primer pair. All primer pairs that overlap with the leading primer pair are scored and sorted, depending on; the size of overlap, number of primers and amplicon size. Each potential primer pair is checked to see if it can be added to any pool other than the leading primer pair’s pool. If a valid primer pair is found, it is added, it becomes the leading, and the process repeats. If not, the solver will attempt to backtrack. This involves removing the leading primer pair and attempting to find a different path. For every possible alternative leading primer pair, the solver tries to find one with a valid overlapping primer pair, allowing the scheme to prevent gaps from being added. However, as the number of possible primer pair combinations is calculated by;

$number\ of\ primer\ pairs^{number\ of\ steps\ back}$ , the number of steps back is limited to 2 for the sake of computational feasibility. If no overlapping primer pairs can be added, a gap is unavoidable, and a walking primer pair is added by iterating over all primer pairs further along the genome. The first primer pair that can be added to any pool is selected and set as the leading primer pair for the next stage.

Scheme mode supports circular genomes, with the `--circular` flag added to another primer pair which spans from the end of the genomes back around to the start, enabling 100% coverage of the genomes after primer trimming.

**Add PrimerPair to the pool**

To add a primer pair into a pool, its primer sequences cannot create any interactions or mispriming PCR products with the primer sequences in the pool. Any interaction with a dimer score less than the threshold (default -26), the primer pair is discarded.

**Interaction checker**

We developed PrimalDimer, a primer dimer detection algorithm based on the work described by Johnston *et al*[3], implemented in the Rust language for performance and optimised for ARTIC PCR conditions. The algorithm takes two primers in opposite orientations and slides them over each other moving one base at a time, at each step the thermodynamic binding stability (ΔG) is calculated via the nearest neighbour parameters, single mismatches and dangling ends[4–6]. Custom penalties and bonuses to account for mismatches, 3' stability, and polymerase binding activity were optimised against a training set of known interacting and non-interacting primers combinations provided by Johnston *et al* [3], expanded to contain previously discovered dimers and non-dimers[7,8]. A ROC analysis was then carried out on a separate testing set of dimers. The output of primaldimer is a score that predicts how severely two primers will interact. A lower score indicates a more impactful dimer.

We ran simulations to determine the maximum number of primers that could be added to a pool before interactions were inevitable (**Supplementary Figure 5)**. For dimer scores -25 to -31, 100 random thermopassing primers are generated, and checked for interactions against a pool of N random primers. The probability was determined from the number of interactions from the 100 random primers. This analysis was repeated 100 times to minimise the effects of randomisation.

**Additional run modes**

Using multiple discrete 3' anchored primers to cover diversity, makes schemes fundamental repairable, as if a unaccounted for mutation is impacting the performance of the cloud, a new primer with the mutation can be rapidly designed and spiked in, ensuring schemes are maintainable and do not require a redesign. Mutations between the primer and the template can be fixed via the `` `repair` `` mode, in which new primer sequences, that account for the mutation, are added into the clouds. If an amplicon is underperforming, the `` `replace` `` mode can generate alternate amplicons, preventing manual laborious iterations. The `interaction` mode enables primer-primer interaction to be profiled in existing schemes.

Visualise primer mismatches. This runmode takes an MSA, containing the primary reference alongside other genomes, and a .bed file. The tool maps primer sequences to their corresponding sites in each genome and calculates the number of mismatches between the primer and the template. This calculation is performed across all primers and input genomes. The resulting data is presented as an interactive heatmap, containing the alignment of primers and corresponding template, facilitating rapid identification of potential primer binding issues across different genetic variants.

**Discrete vs degenerate primers**

The consensus genomes from the measles sequencing were aligned, and used as the input for discrete and degenerate primers. Discrete primers were generated from the internal functions of primalscheme3, degenerate primers used a custom python script to generate the consensus. A 0.25 frequency filter was used to generate degenerate primers only with mutations above the set threshold. Primer Tm are calculated using primer3 with parameters as per `primalscheme3/core/config.py`.

### Supplementary Figures

| Strain | Genotype | ID | NCBI reference |
|---|---|---|---|
| Moraten | A | NCPV 0809211v | |
| Schwartz | A | NCPV 0007233v | AF266291.1 |
| Mvi/Nottingham.GBR/18.04 | D5 | NCPV 0809213v | |
| Mvi/London.UNK/4.02 | D5 | NCPV 0809214v | |
| Mvs/Kingston.GBR/19.06/3 | B3 | NCPV 0809215v | |
| Mvs/London.GBR/25.07 | D4 | NCPV 0809216v | |
| Schwartz | A | NCPV 0809212v | AF266291.1 |
| Edmonston-Zagreb | A | NCPV 0007234v | AY486084.1 |

**Supplementary Table 1.** MeV strain names, genotype, NCPV ID and corresponding NCBI reference where available.

| Clade | ID |
|---|---|
| Clade Ia | KJ642613 |
| Clade Ib | PP601219 |
| Clade IIa | DQ011153 |
| Clade IIb | NC_063383 |

**Supplementary Table 2.** Clade-specific reference genomes used for MPXV scheme remapping.

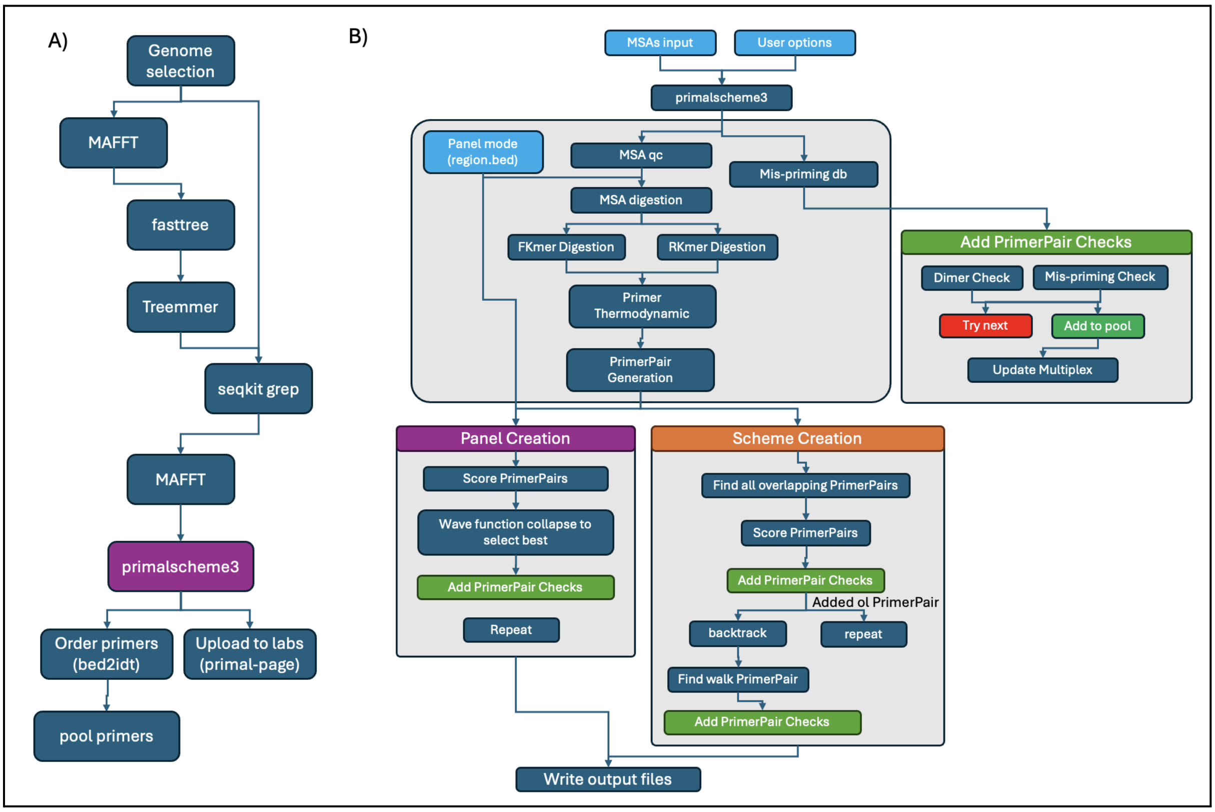


**Supplementary Figure 1. A)** Flow diagram of the preprocessing steps used for preparing input alignment for primalscheme3. **B)** A flow diagram of the main steps performed by primalscheme3, shown separately are the panel and scheme create modes.

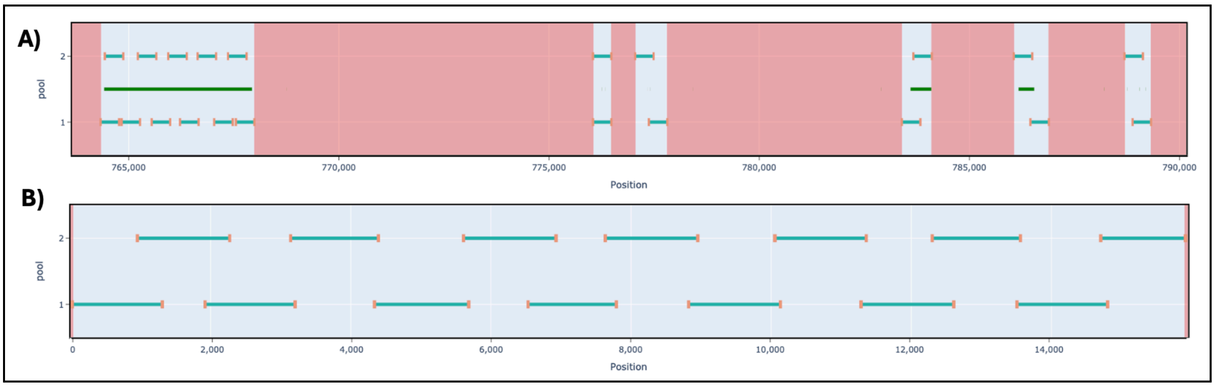


**Supplementary Figure 2 :** Output of the two different solving algorithms. Primer sites are shown in orange, and amplicons in blue. **A)** Panel mode, in which amplicons are added depending on their coverage of desired region (green). **B)** Scheme mode, in which amplicons are progressively added depending on the overlap to the previous amplicon.

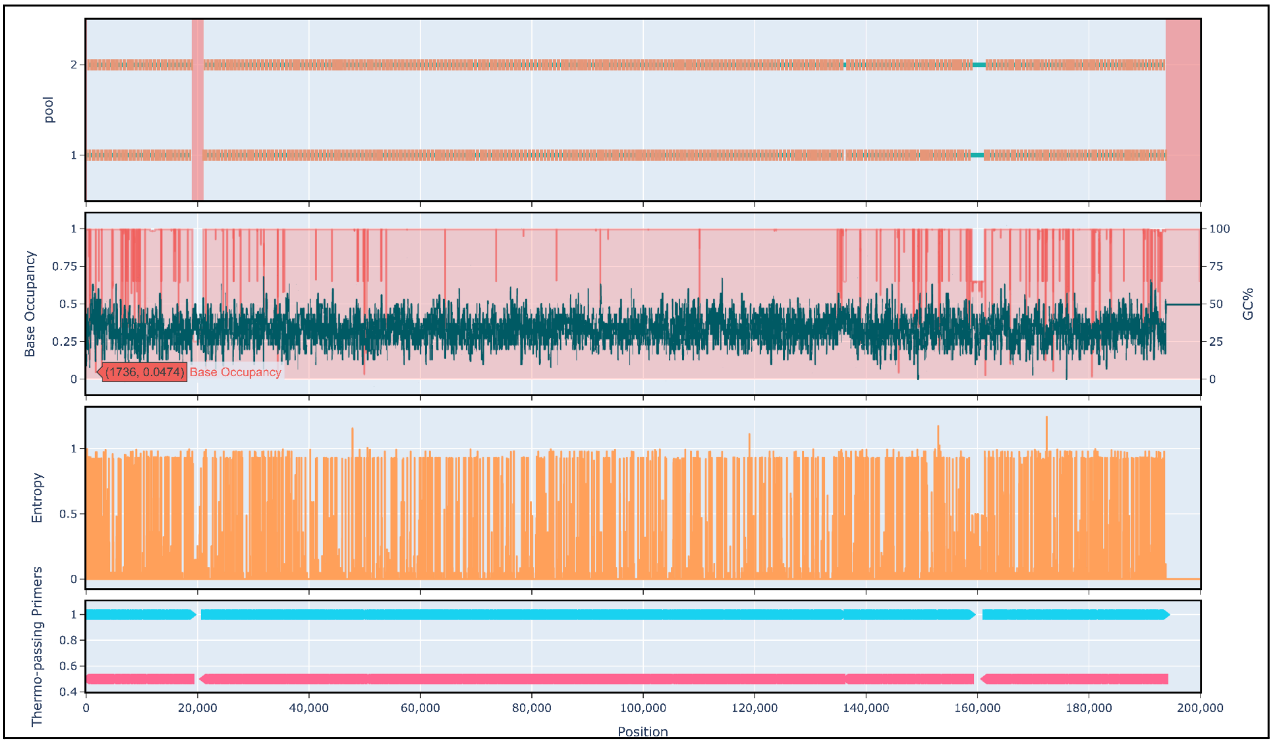


**Supplementary Figure 3:** Screenshot of interactive plot output. The position of the amplicons in the scheme with gaps highlighted in red (top), %GC and base occupancy of the alignment (upper middle), entropy (lower middle), and other possible passing primers (bottom). All positions relative to the primary reference genome.

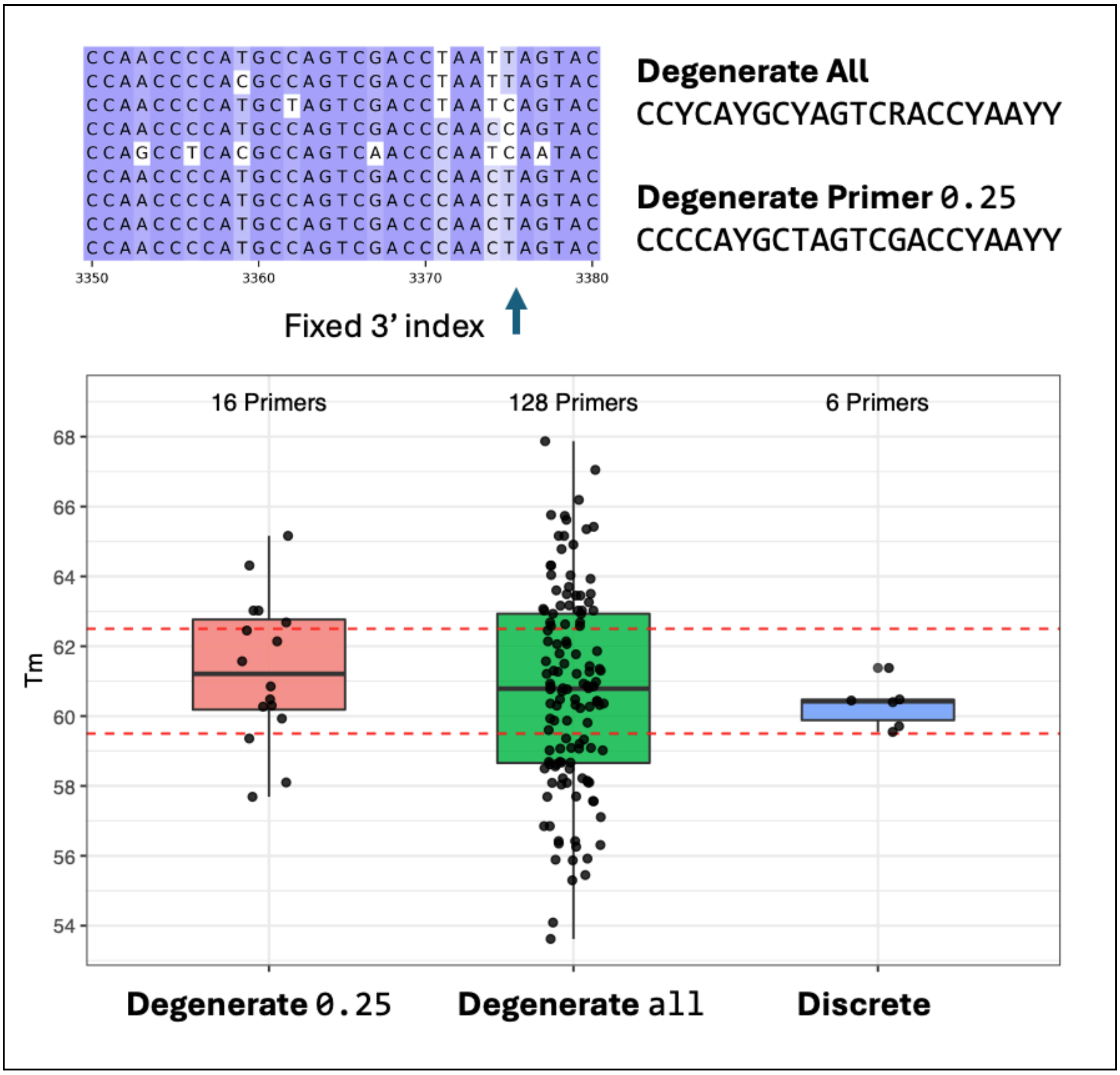


**Supplementary Figure 4:** The number and Tm of primers using degenerate and discrete methods. For a given input alignment (top right) this can be represented as a degenerate sequence using all ambiguous bases or ambiguous bases for variant frequency 0.25 (top right). The Tm (calculated using Primer3) for the product of each degenerate sequence is indicated by a dot with box and whisker plots showing the interquartile range, Tm range used by PrimalScheme indicated by dashed red lines (bottom).

Discrete primers required 6 primers to cover the known diversity in the MSA. The degenerate primers required 16 primers with a Tm range of 7.8 to 65.1, and 128 primers with a Tm range of 55.8 to 67.9 for the 0 and 0.25 frequency filter, respectively.

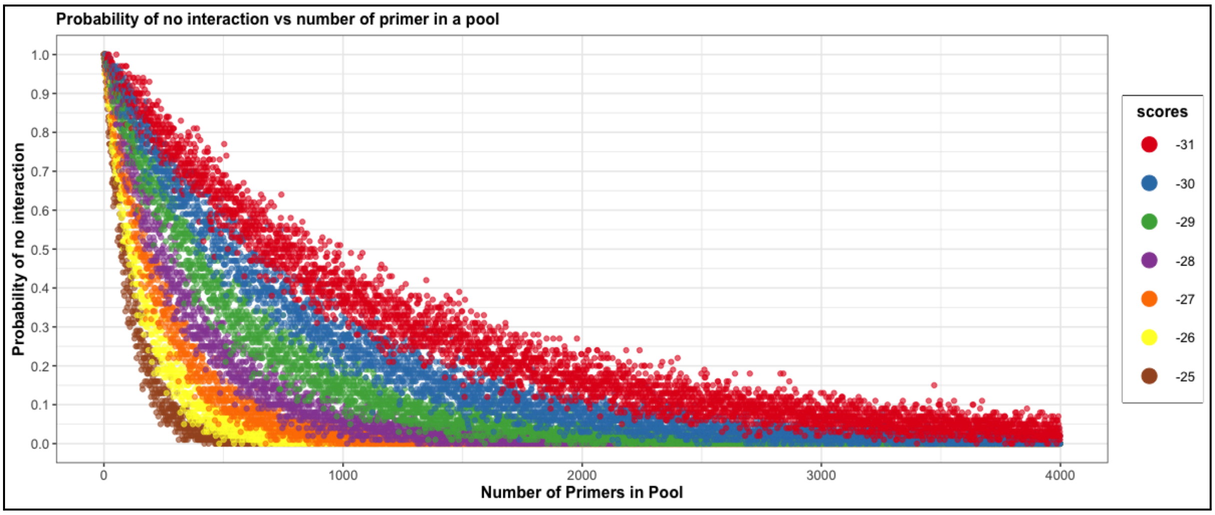


**Supplementary Figure 5**: The finite sequence space limits the number of primers that can be combined into a pool before detection of interactions is unavoidable. With the default dimer score (-26), the probability of a new primer not interacting with a pool containing 1000 primers approaches 0. If the dimer score is laxed to -29, between 2,000 and 3,000 primers can be added before interactions are inevitable. At these scores, the most severe dimers are prevented, however, some dimers will be allowed through.

PrimalDimer performs well at determining primer dimers. Producing an AUC of 0.87 for ROC analysis. The optimal dimer score of '-27' produces a TPR of 0.93 and an FPR of 0.33, and would have prevented all significant dimers found in the early ARTIC SARS-CoV-2 schemes [7], alongside the dimer responsible for the sequencing artefact in v4.1 [8].

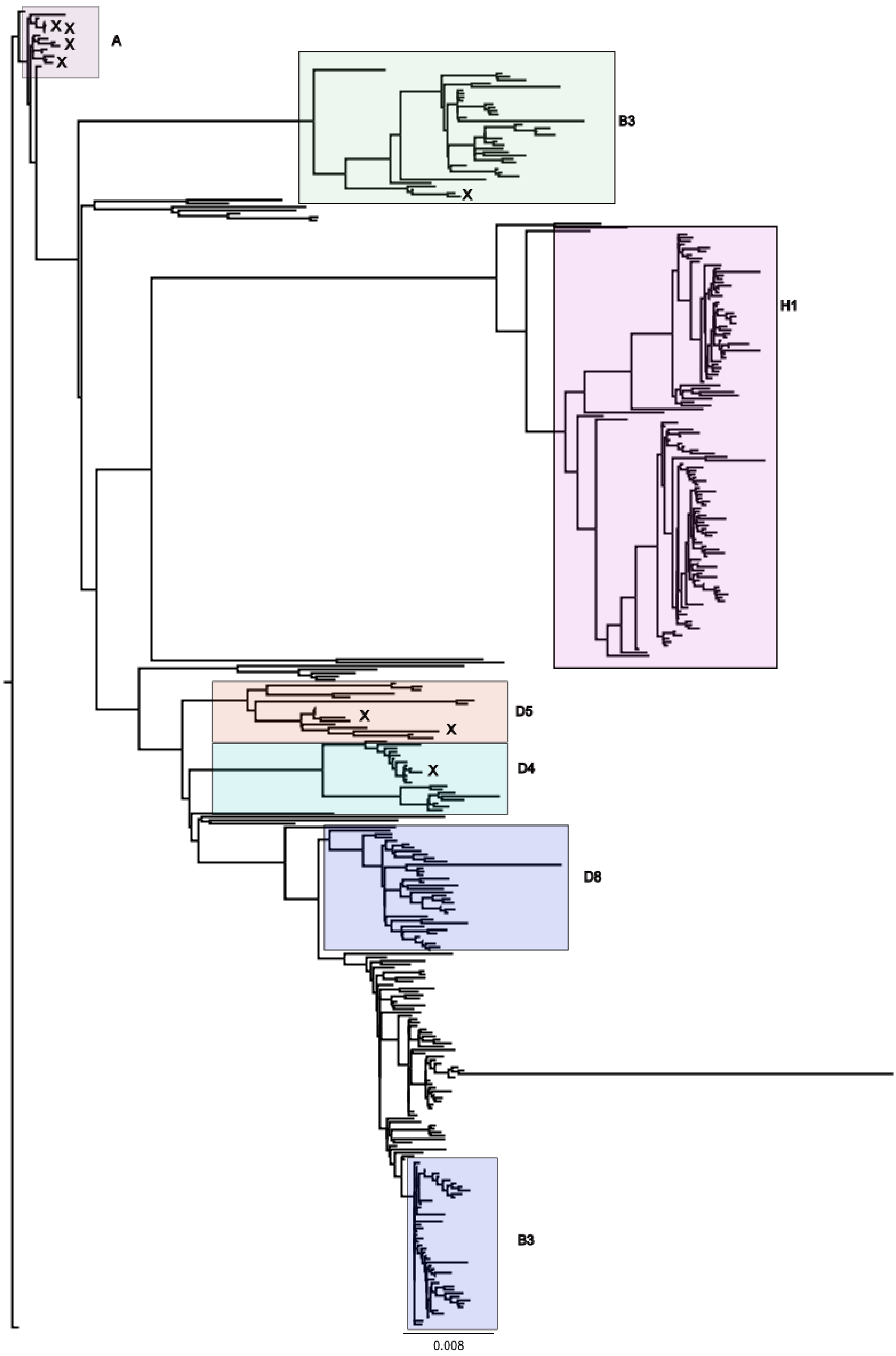
A
X
X
X
X
B3
X
H1
D5
X
X
D4
X
D8
B3
0.008

**Supplementary Figure 6.** Phylogenetic tree generated using FastTree from the input alignment of MeV with consensus sequences for samples added. Clades representing genotypes have been annotated where possible. Positions of consensus sequences are represented by an X.

### Supplementary References


1. Untergasser, A. *et al.* Primer3—new capabilities and interfaces. *Nucleic Acids Res.* **40**, e115–e115 (2012).
2. Quick, J. *et al.* Multiplex PCR method for MinION and Illumina sequencing of Zika and other virus genomes directly from clinical samples. *Nat. Protoc.* **12**, 1261–1276 (2017).
3. Johnston, A. D., Lu, J., Ru, K., Korbie, D. & Trau, M. PrimerROC: accurate condition-independent dimer prediction using ROC analysis. *Sci. Rep.* **9**, 209 (2019).
4. Bommarito, S., Peyret, N. & Jr, J. S. Thermodynamic parameters for DNA sequences with dangling ends. *Nucleic Acids Res.* **28**, 1929–1934 (2000).
5. Peyret, N., Seneviratne, P. A., Allawi, H. T. & SantaLucia, J. Nearest-neighbor thermodynamics and NMR of DNA sequences with internal A.A, C.C, G.G, and T.T mismatches. *Biochemistry* **38**, 3468–3477 (1999).
6. SantaLucia, J. A unified view of polymer, dumbbell, and oligonucleotide DNA nearest-neighbor thermodynamics. *Proc. Natl. Acad. Sci.* **95**, 1460–1465 (1998).
7. Itokawa, K., Sekizuka, T., Hashino, M., Tanaka, R. & Kuroda, M. Disentangling primer interactions improves SARS-CoV-2 genome sequencing by multiplex tiling PCR. *PLOS ONE* **15**, e0239403 (2020).
8. Wilkinson, S., Groves, N., Quick, J. & Loman, N. J. Erroneous Mutations Associated with 64_L-60_R Primer-Dimer in ARTIC 4/4.1 - Laboratory. *ARTIC Real-time Genomic Surveillance* https://community.artic.network/t/erroneous-mutations-associated-with-64-l-60-r-primer-dimer-in-artic-4-4-1/419/1 (2022).
